# A lipid nanoparticle RBD-hFc mRNA vaccine protects hACE2 transgenic mice against lethal SARS-CoV-2 infection

**DOI:** 10.1101/2021.03.29.436639

**Authors:** Uri Elia, Shahar Rotem, Erez Bar-Haim, Srinivas Ramishetti, Gonna Somu Naidu, David Gur, Moshe Aftalion, Ma’ayan Israeli, Adi Bercovich-Kinori, Ron Alcalay, Efi Makdasi, Theodor Chitlaru, Ronit Rosenfeld, Tomer Israely, Sharon Melamed, Inbal Abutbul Ionita, Dganit Danino, Dan Peer, Ofer Cohen

## Abstract

The current global COVID-19 pandemic led to an unprecedented effort to develop effective vaccines against SARS-CoV-2. mRNA vaccines were developed very rapidly during the last year, and became the leading immunization platform against the virus, with highly promising phase-3 results and remarkable efficacy data. Since most animal models are not susceptible to SARS CoV-2 infection, pre-clinical studies are often limited to infection-prone animals such as hamsters and non-human primates. In these animal models, SARS-CoV-2 infection results in viral replication and a mild disease disease. Therefore, the protective efficacy of the vaccine in these animals is commonly evaluated by its ability to elicit immunologic responses, diminish viral replication and prevent weight loss. Our lab recently reported the design of a SARS-CoV-2 human Fc-conjugated receptor-binding domain (RBD-hFc) mRNA vaccine delivered via lipid nanoparticles (LNPs). These experiments demonstrated the development of a robust and specific immunologic response in RBD-hFc mRNA-vaccinated BALB/c mice. In the current study, we evaluated the protective effect of this RBD-hFc mRNA vaccine by employing the K18-hACE2 mouse model. We report that administration of RBD-hFc mRNA vaccine to K18-hACE2 mice led to a robust humoral response comprised of both binding and neutralizing antibodies. In accordance with the recorded immunologic immune response, 70% of vaccinated mice were protected against a lethal dose (3000 plaque forming units) of SARS-CoV-2, while all control animals succumbed to infection. To the best of our knowledge, this is the first non-replicating mRNA vaccine study reporting protection of K18-hACE2 against a lethal SARS-CoV-2 infection.

---

Severe acute respiratory syndrome coronavirus 2 (SARS-CoV-2), identified as the causative agent of coronavirus disease 2019 (COVID-19), developed in the last year into a global pandemic, causing (as of March 20^th^, 2021) over 120 million cases and 2.7 million deaths worldwide.^1^ Unprecedented international collaboration has led to record time development of several vaccine candidates, of which the mRNA vaccine platform has proven to be highly efficacious in phase 3 clinical studies.^2,3^ Consequently, mRNA-based vaccines became the first prophylactic therapeutics to be approved by the UK’s Medicine and Healthcare products Regulatory Agency (MHRA), and the US Food and Drug Administration (FDA), authorizing the Pfizer-BioNTech BNT162b2 vaccine, followed by approval of Moderna’s mRNA-1273 candidate for global emergency mass use. The high efficacy of mRNA vaccines in eliciting a robust and effective immunologic response, together with high safety profile and simple, rapid manufacturing process, make this platform extremely attractive for the prevention of infectious diseases.

The main obstacle in the *in vivo* administration of mRNA is its efficient delivery to target cells. In recent years, a wide range of different delivery approaches were developed, mainly in the context of *in vivo* siRNA delivery, some of which have been implemented also for the delivery of mRNA. Lipid nanoparticles (LNPs) represent the most advanced and frequently used carrier platform for mRNA delivery, and is currently the standard method being used to introduce mRNA vaccines to humans participating in clinical trials worldwide. LNPs are generally comprised of a combination of four elements: cholesterol, a phospholipid, polyethylene glycol (PEG)-linked lipid and an ionizable lipid. Of these components, the ionizable lipid has been shown to play a central role in the effective delivery and subsequent translation of the mRNA and encoded-protein expression.^4,5^ In the current study, we directly examined the *in vivo* efficacy of an LNP-encapsulated human Fc-conjugated SARS-CoV-2 receptor binding domain (RBD-hFc)-based mRNA vaccine encompassing an ionizable lipid (lipid #14) previously demonstrated to be highly potent in eliciting a robust immune response in vaccinated BALB/c mice.^6^ In order to evaluate the efficacy of the vaccine against viral infection, the human ACE2 (K18-hACE2) transgenic mice model was employed, a model previously shown to recapitulate the susceptibility to SARS-CoV-2 infection. Prime-boost intramuscular immunization of K18-hACE2 mice with 5 μg LNP RBD-hFc mRNA led to 70% protection against an lethal 3*10^3^ plaque forming units (PFU) SARS-CoV-2 intranasal infection which resulted in the death of all untreated animals. To the best of our knowledge, this is the first report demonstrating high efficacy of a conventional (non-replicating) mRNA vaccine against lethal SARS-CoV-2 infection in a hACE2 mice model.

An LNP RBD-hFc mRNA vaccine was designed, based on a previous lipid formulation screen that we recently reported.^6^ The results of the screen led to the selection of lipid #14, which was highly potent in eliciting strong humoral and cellular responses in BALB/c mice immunization studies. A schematic representation of the hFc-fused RBD mRNA construct is shown in Figure 1a. Following LNP preparation, samples were analyzed for size and uniformity by dynamic light scattering (DLS). As can be seen in Figure 1b, particles were small (∼55 nm) and uniformly distributed, as evidenced by an average polydispersity index (PDI) of <0.2. Cryogenic electron microscopy (Cryo-EM) analysis supported the DLS data, showing small and uniform particles (Figure 1c).

**Figure 1.**
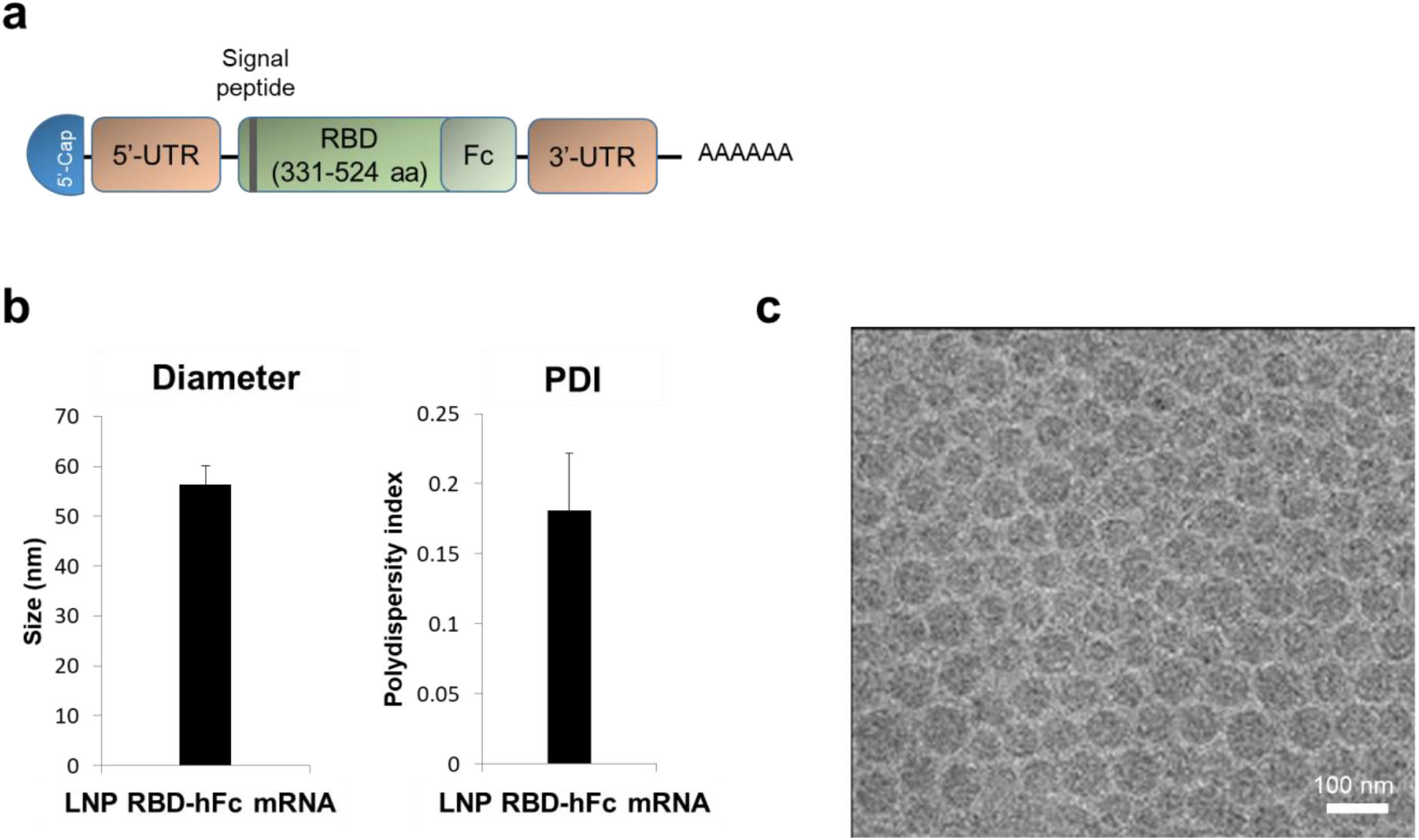
Design of LNP RBD-hFc mRNA and physicochemical characterization. (a) Schematic representation of the RBD-hFc mRNA construct. (b) Size distribution and polydispersity index (PDI) of LNPs measured by dynamic light scattering. (c) A representative Cryo-EM image of LNP-encapsulated RBD-hFc mRNA. Scale bar 100nm.

In order to evaluate the translation of RBD-hFc mRNA into a functional protein, we performed *in vitro* transfection of HEK293 cells with LNP RBD-hFc mRNA, and determined protein expression and target binding. Western blot analysis established expression of RBD-hFc in the supernatant but not in the cell pellets, confirming the extracellular secretion of the translated RBD-hFc exhibiting the expected molecular size (Figure 2a). To confirm the functionality of the translated RBD-hFc, we applied ELISA against hACE2. Binding was evaluated for cell supernatant samples collected 72 h following transfection with LNPs-encapsulated RBD-hFc mRNA, empty LNPs or RBD-hFc-expression plasmid. As demonstrated (Figure 2b), comparable binding was observed for RBD-hFc expressed by both mRNA and control DNA plasmid, but not for the empty LNPs.

**Figure 2.**
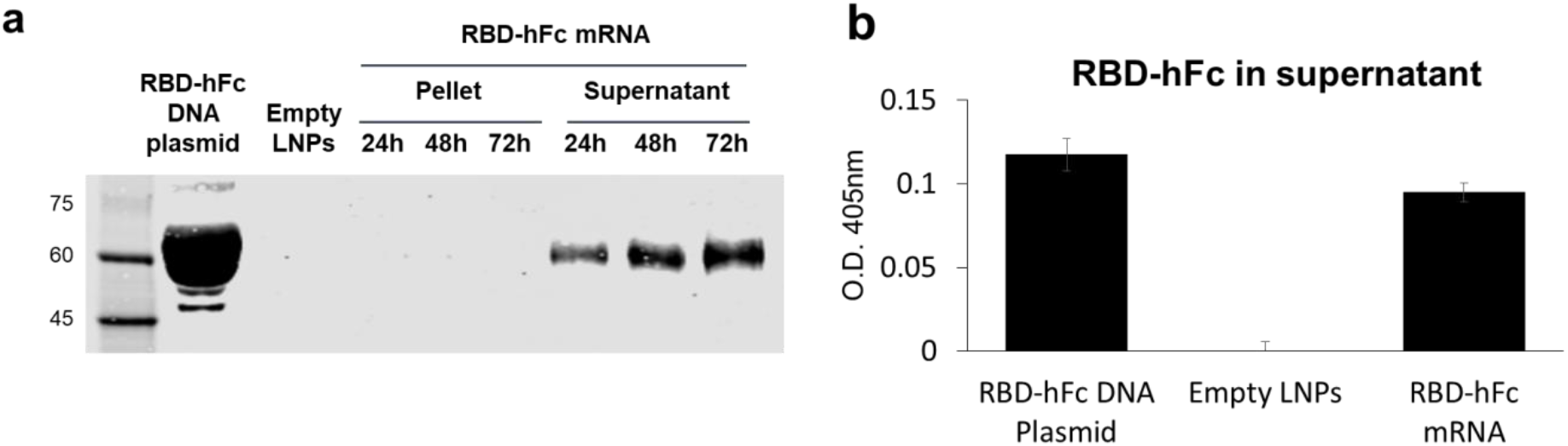
*In vitro* charaterization of RBD-hFc expression. (a) Western blot analysis of HEK293 cells transfected with RBD-hFc DNA plasmid (1.5 μg/ml, 72 h), empty LNPs (72 h) or LNP RBD-hFc mRNA (1 μg/ml, 24-72 h). Anti-hACE2 ELISA for the evaluation of hACE2 binding by hFc-RBD. Plates were coated with hACE2 (2 μg/ml), and supernatant fractions of RBD-hFc DNA plasmid-, empty LNPs- or LNP RBD-hFc mRNA-transfected cells (72 h) were analyzed.

ACE2 receptor is the main mediator of SARS-CoV-2 binding and entry into infected cells. However, mouse ACE2 does not support binding of SARS-CoV-2, and therefore conventional laboratory mice strains are not suitable for infection studies.^7^ In order to evaluate the *in vivo* efficacy of the LNP RBD-hFc mRNA vaccine against a lethal SARS-CoV-2 challenge, we employed a human ACE2 (K18-hACE2) transgenic mice model, in which hACE2 expression is driven by the epithelial cell cytokeratin-18 (K18) promoter. This model has been shown to enable efficient SARS-CoV-2 infection, leading to severe disease and lethality, and has been the model of choice for several SARS-CoV-2 immunization and therapy studies.^8-10^ Groups of 6-8 weeks old K18-hACE2 mice were intramuscularly administered with 5 μg LNP RBD-hFc mRNA. Control experimental groups were immunized with empty LNPs or recombinant RBD-hFc (rRBD) protein (10 μg, subcutaneously). Prime-only or prime-boost vaccination regimens were employed, with the animals being primed at day 0 and boosted 23 days later (see vaccination outline in Figure 3a). To characterize the humoral response of the vaccinated animals, serum samples were collected prior to challenge.

**Figure 3.**
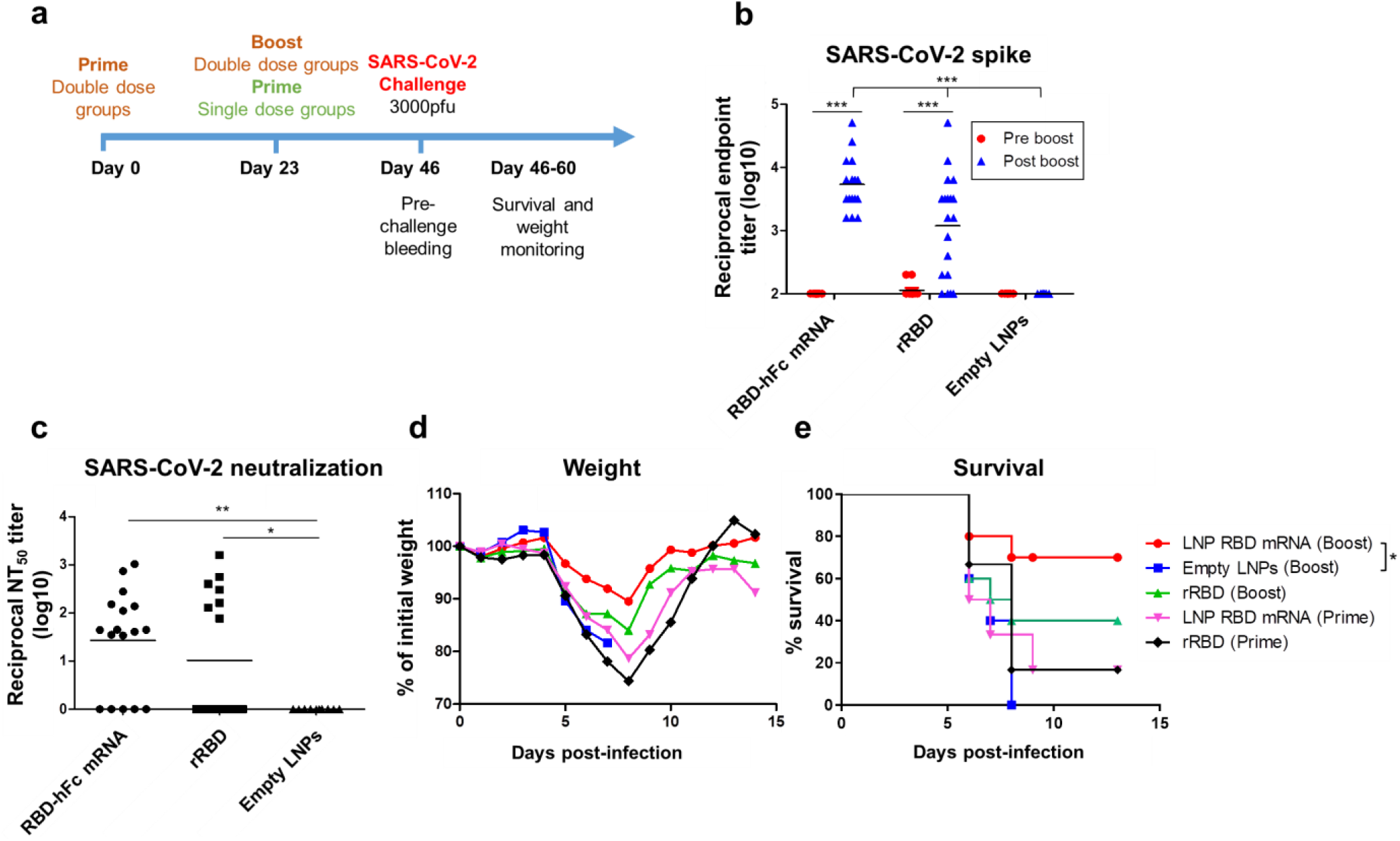
LNP RBD-hFc mRNA vaccine protects hACE2 trangenic mice against a lethal SARS-CoV-2 challenge. (a) Schematic diagram of immunization regimens and serum samples collection. K18-hACE2 mice were intramuscularly administered with LNP RBD-hFc mRNA (5 μg) or rRBD (10 μg) at a prime-only (*n=6*) or prime-boost (*n=10*) vaccination regimen. Control animals were administered with empty LNPs (prime-boost) (*n=5*). Serum samples were collected 23 (“pre boost”) and 46 (“post boost”) days after priming, and were assayed for SARS-CoV-2 spike-specific IgG antibodies (b) and neutralizing antibodies against SARS-CoV-2 (c) as described in the methods section. Animals were then intranasally infected with 3*10^3^ PFU of SARS-CoV-2, and monitored for weight loss (d) and survival (e). Statistical analysis was performed using a two-way ANOVA with Bonferroni’s multiple comparisons test (for ELISA data), one-way ANOVA followed by post hoc Newman– Keuls test (for neutralizing antibodies data) or log-rank (Mantel–Cox) test (for survival data) (*, p < 0.05; **, p < 0.01; ***, p < 0.001).

As shown in Figure 3b, a single intramuscular immunization (“pre-boost”) with either LNP RBD-hFc mRNA (n=10) or rRBD (n=11), did not elicit a significant humoral response, with most animals exhibiting anti-spike titers of <100. Conversely, a robust and statistically significant antibody response was observed in both LNP RBD-hFc mRNA (n=16) and rRBD (n=19) prime-boost immunization groups (“post-boost”) compared with the empty LNPs control group (n=8). Within the immunization groups, the recorded anti-spike titers were significantly higher in mice administered with LNP RBD-hFc mRNA (mean log_10_ titer=3.7) compared with rRBD-administered mice (mean log_10_ titer =3.0). We next evaluated the post-boost *in vitro* neutralizing antibody response using a SARS-CoV-2 plaque reduction neutralization test (PRNT). In line with the binding antibody titers, a substantial neutralizing antibody response was detected in both immunization groups, with the LNP RBD-hFc mRNA group exhibiting a stronger response compared with the rRBD group (although not statistically significant) (Figure 3c). Twenty-three days following the last administered dose, all mice were challenged with 3*10^3^ PFU of SARS-CoV-2 *via* the intranasal route, and monitored daily for weight loss and survival. As shown in figure 3d, weight loss was recorded in all animal groups, starting at day 5 and reaching a maximal loss at day 8 post-challenge. Thereafter, surviving animals began to steadily recover and return to initial weight approximately at day 11-12 post-challenge. Mice receiving empty LNPs, or single administration of either LNP RBD-hFc mRNA or rRBD, showed significant drop in weight (∼20-25% weight loss). In contrast, mice administered with a booster dose exhibited a less pronounced weight loss of 10% and 15% for LNP RBD-hFc mRNA and rRBD immunization groups, respectively (Figure 3d). Most importantly, while the 3*10^3^ PFU SARS-CoV-2 challenge was 100% (5/5) fatal to empty LNPs-administered control animals, prime-boost vaccination of mice with LNP RBD-hFc mRNA resulted in a 70% (7/10) survival rate. In accordance with the recorded humoral response, which was more robust in the LNP RBD-hFc mRNA group, prime-boost vaccination with rRBD led to a lower survival rate of 40% (4/10). Lastly, supported by the low antibody titers recorded pre-boost, only 16% (1/6) of mice receiving prime-only vaccinations survived (Figure 3e).

mRNA vaccines against SARS-CoV-2 have been studied extensively in the past year, both in pre-clinical and clinical studies.^11^ Most SARS-CoV-2 mRNA vaccine pre-clinical studies begin with evaluation of the immunologic response elicited by the vaccine in animal models which are not susceptible to SARS-CoV-2 infection (i.e. BALB/c, C57/BL6 mice). In order to examine the efficacy of the vaccine, SARS-CoV-2 infection-prone animal models are frequently used (e.g. hamsters and non-human primates). In many studies, these animals are vaccinated and then challenged with virulent SARS-CoV-2, leading to symptomatic disease, weight loss and extensive viral replication in different tissues, but not death. Therefore, the protective effect of the vaccine is commonly evaluated by its ability to diminish viral replication, elicit a robust immune response and reduce weight loss. ^12-14^

In the current study, we sought to evaluate the efficacy of the vaccine by using the K18-hACE2 mouse model, which has been shown to be highly susceptible to SARS-CoV-2 infection. Prime-boost immunization of K18-hACE2 mice with LNP RBD-hFc mRNA vaccine, elicited a specific humoral response, as evidenced by the detection of binding-as well as neutralizing antibodies. Most importantly, administration of a lethal dose of 3*10^3^ PFU SARS-CoV-2 led to seventy percent survival in immunized mice, compared with full mortality in the control group.

Only a few studies have evaluated so far the efficacy of SARS-CoV-2 mRNA vaccines in hACE2 transgenic mice. A recent preprint study by Arcturus and Singapore’s Duke-NUS Medical School consortium described full protection of K18-hACE2 mice following a single dose of self-transcribing and replicating RNA (STARR™)-based vaccine candidate.^15^ A different study by the Chinese Academy of Sciences also employed a hACE2 mouse model for evaluation of vaccine protection efficacy.^16^ The data in that study, however, is based on immune humoral response, weight loss, lung viral titer and lung immunohistochemistry analysis, and not on survival of immunized animals. In summary, to the best of our knowledge, the current study demonstrates for the first time SARS-CoV-2 mRNA vaccine-mediated protection of K18-hACE2 mice against an otherwise lethal viral infection.

## Acknowledgments

We thank Hila Cohen, Liat Bar-On, Shirley Lazar, Tal Noy-Porat, and Yinon Levi for their assistance and support and Shmuel Yitzhaki, Ohad Mazor, and Emanuelle Mamroud for their insightful discussions and encouragement. We also thank Amir Rosner, Tseela David and Beni Shareabi for animal husbandry.

We thank Inbar Freilich for her professional contribution to the Cryo-EM analysis.

## Conflict of interest

The authors declare the following competing financial interest(s): D.P. receives licensing fees (to patents on which he was an inventor) from, invested in, consults (or on scientific advisory boards or boards of directors) for, lectured (and received a fee) or conducts sponsored research at TAU for the following entities: Alnylam Pharmaceuticals Inc., Arix Biosciences Inc., ART Biosciences, BioNtech RNA Pharmaceuticals, Centricus, Diagnostear Ltd., EPM Inc., Earli Inc., lmpetis Biosciences, Kernal Biologics, GPCR Inc., Medison Pharma Ltd., Newphase Ltd., NLC Pharma Ltd., Nanocell Therapeutics, NanoGhosts Ltd., Precision Nanosystems Inc., Paul Hastings Inc., Regulon, Roche, SciCann, Shire Inc., SirTLabs Corporation, VLX Ventures, TATA Cooperation, Teva Pharmaceuticals Inc., and Wize Pharma Ltd. All other authors declare no competing financial interests.

## Materials and methods

### Cells and animals

HEK293T cells (ATCC CRL-3216) and Vero E6 cells (ATCC CRL-1586) were maintained at 37°C, 5% CO_2_ in Dulbecco’s modified Eagle’s medium (DMEM) supplemented with 10% fetal bovine serum (FBS), MEM non-essential amino acids (NEAA), 2 mM L-glutamine, 100 Units/ml penicillin, 0.1 mg/ml streptomycin and 12.5 Units/ml Nystatin (P/S/N) (Biological Industries, Israel).

Female K18-hACE2 (B6.CgTg(K18ACE2)2Prlmn/J HEMI) mice (6–8 weeks old) were obtained from Jackson and randomly assigned into cages in groups of 10 animals. The mice were allowed free access to water and rodent diet (Harlan, Israel). All animal experiments were conducted in accordance with the guideline of the Israel Institute for Biological Research (IIBR) animal experiments committee: protocol numbers M-66-20.

### mRNA

CleanCap, pseudouridine-substituted Fc-conjugated RBD mRNA (amino acids 331– 524; GenBank: QHD43416.1) was purchased from TriLink Bio Technologies (San Diego, CA, USA). The Fc-conjugated RBD mRNA was designed to include the exact translated RBD-hFc protein sequence as the recombinant protein.

### LNP Preparation and Characterization

LNPs were synthesized by mixing one volume of lipid mixture of ionizable lipid, DSPC, cholesterol, and DMG-PEG (40:10.5:47.5:2 mol ratio) in ethanol and three volumes of mRNA (1:23 (w/w) mRNA to lipid) in acetate buffer. Lipids and mRNA were injected in to a microfluidic mixing device Nanoassemblr (Precision Nanosystems, Vancouver, BC, Canada) at a combined flow rate of 12 mL/min. The resultant mixture was dialyzed against phosphate buffered saline (PBS; pH 7.4) for 16 h to remove ethanol.

Particles in PBS were analyzed for size and uniformity by dynamic light scattering (DLS, Malvern Instruments Ltd., Worcestershire, U.K.). RNA encapsulation in LNPs was calculated according to Quant-iT RiboGreen RNA Assay Kit (Thermo Fisher, Waltham, MA, USA).

### Cryogenic Electron Microscopy (cryo-EM)

Samples were prepared in a closed chamber at a controlled temperature and at water saturation. A 5-6 µl drop of each suspension was placed on a 200-mesh transmission electron microscope (TEM) copper grid covered with a perforated carbon film. The drop was blotted, and the sample was plunged into liquid ethane (−183°C) to form a vitrified specimen, then transferred to liquid nitrogen (–196°C) for storage. Vitrified specimens were examined at temperatures below −175°C in a Talos F200C with field emission gun operated at 200Kv or a Tecnai T12 G2 TEM (FEI, Netherlands). Images were recorded on a Cooled Falcon IIIEC (FEI) Direct Detection Device by TIA software attached to the Talos or a Gatan MultiScan 791 camera by DigitalMicrograph software (Gatan, U.K.) on the Tecnai. Volta PhasePlate (VPP) was used for contrast enhancement. Images are recorded in the low-dose imaging mode to minimize beam exposure and electron-beam radiation damage using lab procedures.^17,18^

### *In vitro* transfection

LNPs-encapsulated RBD-hFc mRNA (1 μg/ml) or empty LNPs were added to 3*10^5^ HEK293T cells seeded in a 6-well plate with 2ml DMEM supplemented with 10% fetal bovine serum (FBS). For positive control, cells were transfected with RBD-hFc-expressing pcDNA3 vector (1.5 μg/ml) using jetPEI (Polyplus-transfection) according to manufacturer’s protocol.

### Western blot

In order to evaluate RBD-hFc expression in transfected cells, supernatants were collected, and cells were lysed on ice in RIPA buffer (Merck) at 24, 48 or 72 h after transfection. Lysates were nutated at 4°C for 10 min, then centrifuged at 20,000×g for 10 min at 4°C. RBD-hFc was purified from whole-cell lysates and supernatants using Protein A HP SpinTrap (GE) and eluted with 100ul IgG elution buffer (Pierce). Samples were then separated by 4–12% polyacrylamide Bis-tris gel electrophoresis (Invitrogen) and blotted onto a nitrocellulose membrane. The membrane was blocked for 1 h in Intercept® Blocking Buffers (Li-COR). RBD-hFc was detected by incubation of the membrane with purified IgG fraction from serum of rabbit immunized with SARS-CoV-2 spike protein for over-night at 4°C, followed by a secondary antibody IRDye 680RD goat anti-rabbit (LIC-92668071) incubation of 1 h at RT. Reactive bands were detected by Odyssey CLx infrared imaging system (LI-COR).

### Production of recombinant SARS-CoV-2 proteins

Recombinant SARS-CoV-2 spike glycoprotein for ELISA and human Fc-RBD-fused protein for vaccination were designed and expressed as previously described.^19^ Briefly, a stabilized soluble version of the spike protein (based on GenPept:QHD43416 ORF amino acids 1–1207) was produced using an ExpiCHO Expression system (Thermoscientific, USA, Cat. No. A29133) following purification using HisTrap (GE Healthcare, U.K.) and Strep-TactinXT (IBA, Germany). The purified protein was sterile-filtered and stored in PBS.

Human Fc-RBD-fused protein was expressed using previously designed Fc-fused protein expression vector,^20^ giving rise to a protein comprising two RBD moieties (amino acids 331–524; see accession number of the S protein above) owing to the homodimeric human (γ1) Fc domain (huFc). Expression of the recombinant proteins was performed using the ExpiCHO Expression system (Thermoscientific) following purification using HiTrap Protein-A column (GE Healthcare). The purified protein was sterile-filtered and stored in PBS.

### Animal Vaccination Experiments

For RBD-hFc mRNA vaccination studies, groups of 6–8 week old female K18-hACE2 mice were administered intramuscularly (50 μL in both hind legs) with SARS-CoV-2 RBD-hFc mRNA (5 μg) encapsulated with LNP formulation #14. For recombinant RBD-hFc vaccination studies, mice were administered subcutaneously (100 μL) with rRBD (10 μg). Both RBD-hFc mRNA- and rRBD -immunized animals were either prime-only administered, or boosted at day 23 with the same priming dose administered on day 0. Control animals were administered with empty LNPs (prime-boost). Serum samples were collected on day 23 (“ pre boost”) and 46 (“ post boost”) for evaluation of SARS-CoV-2 spike-specific humoral response.

### SARS-CoV-2 infection

All animal experiments involving SARS-CoV-2 were conducted in a BSL3 facility.

Infection experiments were carried out using SARS-CoV-2, isolate Human 2019-nCoV ex China strain BavPat1/2020 that was kindly provided by Prof. Dr. Christian Drosten (Charité, Berlin, Germany) through the European Virus Archive—Global (EVAg Ref-SKU: 026V-03883), as recently described. ^9^

All experimental animal groups (prime-only RBD-hFc mRNA, prime-only rRBD, prime-boost RBD-hFc mRNA, prime-boost rRBD and prime-boost empty LNPs) were anesthetized and intranasally inoculated with 3*10^3^ PFU of SARS-CoV-2 BavPat1/2020 virus diluted in PBS supplemented with 2% FBS (Biological Industries, Israel). Animal body weight was monitored daily throughout the follow-up period post-infection. Mice were evaluated once a day for clinical signs of disease and dehydration. Euthanasia was applied when the animals exhibited irreversible veterinary-evaluated disease symptoms (rigidity, lack of any visible reaction to contact).

### ELISA

Direct ELISA was performed for the detection of SARS-CoV-2 spike-specific antibodies in immunized mouse sera. MaxiSORP ELISA plates (Nunc) were pre-coated with spike protein (2 μg/mL) in NaHCO_3_ buffer (50 mM, pH 9.6) overnight. Plates were washed three times and blocked with 2% BSA (Sigma-Aldrich, #A8022) in PBST for 1 h at 37°C. After three washes with PBST, plates were incubated with serially diluted mouse sera in PBST/BSA for 1 h at 37°C. Following washing, goat anti-mouse alkaline phosphatase-conjugated IgG (Jackson Immuno Research Laboratories, No. 115-055-003) was added for 1 h at 37°C. The plates were washed with PBST and reactions were developed with *p*-nitrophenyl phosphate substrate (PNPP; Sigma-Aldrich, N2765). Plates were read at 405 nm absorbance, and antibody titers were calculated as the highest serum dilution with an OD value above 2 times the average OD of the negative controls.

Evaluation of *in vitro* RBD-hFc expression in transfected HEK293 cells was performed essentially as described above. Briefly, Plates were pre-coated with recombinant hACE2 (2 μg/mL) overnight, then incubated with whole cell lysates and pellets of transfected cells, followed by AP-conjugated Donkey anti-human IgG (Jackson Immuno Research Laboratories, No. 709-055-149). Detection was performed using the PNPP substrate.

### Plaque reduction neutralization test (PRNT)

Handling of SARS-CoV-2 was conducted in a BSL3 facility in accordance with the biosafety guidelines of the Israel Institute for Biological Research (IIBR). SARS-CoV-2 (GISAID accession EPI_ISL_406862) strain was kindly provided by Bundeswehr Institute of Microbiology, Munich, Germany. Virus stocks were propagated and tittered by infection of Vero E6 cells as recently described.^21^ For PRNT, Vero E6 cells were plated overnight (as detailed above) at a density of 5 × 10^5^ cells/well in 12-well plates. Serum samples were tenfold serially diluted in 200μl of MEM supplemented with 2% FBS, NEAA, 2 mM L-glutamine and P/S/N. Two hundred microliters containing 500 PFU/ml of SARS-CoV-2 virus were then added to the diluted serum solution and the mixture was incubated at 37°C, 5% CO_2_ for 1 h. Monolayers were then washed once with DMEM w/o FBS and 200 μl of each serum-virus mixture was added in triplicates to the cells for 1 h. Virus mixture w/o serum was used as control. In all, 2 ml/well overlay [MEM containing 2% FBS and 0.4% Tragacanth (Sigma, Israel)] was added to each well and plates were further incubated at 37°C, 5% CO_2_ for 48 h. Following incubation, the overlay was aspirated and the cells were fixed and stained with 1 ml of crystal violet solution (Biological industries, Israel). The number of plaques in each well was scored and the NT_50_ (Serum dilution at which the plaque number was reduced by 50%, compared to plaque number of the control, in the absence of serum) was calculated using the Prism software version 8 (GraphPad Software Inc., USA).

### Statistical Analysis

All values are presented as mean plus standard error of the mean (SEM). Statistical analysis was performed using a two-way ANOVA with Bonferroni’s multiple comparisons test (for ELISA data), one-way ANOVA followed by post hoc Newman–Keuls test (for neutralizing antibodies data) or log-rank (Mantel–Cox) test (for survival data) (*, p < 0.05; **, p < 0.01; ***, p < 0.001). All statistical analyses were performed using GraphPad Prism 8 statistical software.

